# Human Skeletal Muscle has Large Capacity to Increase Carnosine Content in Response to Beta-Alanine Supplementation. *A Systematic Review with Bayesian Individual and Aggregate Data E-Max Model and Meta-Analysis*

**DOI:** 10.1101/870584

**Authors:** Nathalia Saffioti Rezende, Paul Swinton, Luana Farias de Oliveira, Rafa Pires da Silva, Vinicius Eira da Silva, Kleiner Nemezio, Guilherme Yamaguchi, Guilherme G Artioli, Bruno Gualano, Bryan Saunders, Eimear Dolan

## Abstract

Beta-alanine (BA) supplementation increases muscle carnosine content (MCarn), and is ergogenic in many situations. Currently, many questions on the nature of the Mcarn response to supplementation are open, and the response to these has considerable potential to enhance the efficacy and applications of this supplementation strategy.

**Objective:** To conduct a Bayesian analysis of available data on the Mcarn response to BA supplementation.

**Methods:** A systematic review with meta-analysis of individual and published aggregate data using a dose response (Emax) model was conducted. The protocol was designed according to PRISMA guidelines. A three-step screening strategy was undertaken to identify studies that measured the Mcarn response to BA supplementation. In addition, individual data from 5 separate studies conducted in the authors’ laboratory were analysed. Data were extracted from all controlled and uncontrolled supplementation studies conducted on healthy humans. Meta-regression was used to consider the influence of potential moderators (including dose, sex, age, baseline Mcarn and analysis method used) on the primary outcome.

**Results and Conclusion:** The Emax model indicated that human skeletal muscle has large capacity for non-linear Mcarn accumulation, and that commonly used BA supplementation protocols may not come close to saturating muscle carnosine content. Neither baseline values, nor sex, appear to influence subsequent response to supplementation. Analysis of individual data indicated that Mcarn is relatively stable in the absence of intervention, and effectually all participants respond to BA supplementation (99.3% response [95%CrI: 96.2 – 100]).

## INTRODUCTION

Carnosine is a dipeptide formed from the amino acids β-alanine and L-histidine, and is present in high concentrations in human skeletal muscle (approximately 20 - 30 mmol·kg^-1^ dry muscle). Its purported roles include: proton buffering [1]; anti-oxidation [2]; anti-glycation; metal chelation [3] and influencing calcium sensitivity [4,5], and hence muscle contractility. Although *in vitro* evidence supports carnosine’s capacity to contribute to each of these processes, the strongest line of *in vivo* and *in vitro* evidence supports an important role for carnosine in intracellular skeletal muscle buffering [3,6]. The pKa of carnosine’s imidazole ring (6.83 [7]) renders it ideally placed to aid in pH regulation within the physiological range of skeletal muscle (which decreases from approximately 7.1 at rest to 6.5 after exhaustive exercise [8]). This mechanistic action is particularly relevant in a sporting context, given that sustained high-intensity efforts are largely fuelled by anaerobic bioenergetic pathways, which lead to hydrogen ion accumulation and an acidic environment. Acidosis is known to contribute to fatigue and limit performance via a wide range of mechanisms [9]. As such, the presence of intracellular pH buffers such as carnosine are essential to maintain high intensity muscle contraction, and hence to sustain performance.

Given the importance of carnosine to athletic performance, considerable research efforts have been made to investigate both means to increase it, and in what situations such increases are ergogenic. In 2006, Harris and colleagues [10] published a series of studies indicating that β-alanine (BA) availability was the limiting factor in intramuscular carnosine synthesis, and that supplementation with this amino acid could substantially increase muscle carnosine content (MCarn). Shortly after, the same group reported that BA supplementation (mean of 5.2 g·day^-1^ for 4 weeks) was ergogenic to high-intensity exercise performance [11]. Since then, the ergogenic benefits of this supplementation strategy have been tested and proven in a wide range of models and recently, our group published a meta-analysis showing that BA supplementation is most ergogenic in capacity-based exercise tests that last between 30 seconds and 10 minutes [12]. This strong evidence supporting the ergogenic potential of BA supplementation has earned it its place as one of the world’s most popular sports supplements, and it is one of just five ergogenic supplements endorsed by the International Olympic Committee [13].

It seems that substantial amounts of BA are required to increase MCarn content, with most studies using doses of approximately 3.2 – 6.4 g·day^-1^, for periods ranging from 4 – 24 weeks. But many questions about the nature of the muscle carnosine response to BA supplementation remain open, and substantial research efforts are being made to refine BA dosing strategies in order to optimize its efficacy and applicability [14]. For example, inter-individual variation in response to supplementation is high [12], yet little is known about what factors underpin this [15]. What is the capacity of the muscle to uptake BA and increase MCarn? What is the individual proportion of response to BA supplementation, and do baseline levels dictate the extent of this? Do sex or age influence response to supplementation? To address these questions, we employed a comprehensive analysis that included individual participant data from multiple studies conducted within our laboratory; and combined these findings with summary published data using a frequently used dose-response model (Emax).

## METHODS

The protocol for this study was designed according to the Preferred Reporting Items for Systematic Reviews and Meta-Analysis (PRISMA) guidelines. The *Population, Intervention, Comparator, Outcomes and Study Design (PICOS)* approach was used to guide the inclusion and exclusion of studies for this review and are described in Table 1.

**Table 1:** Study Inclusion and Exclusion Criteria

|  |  |
| --- | --- |
| **Population:** | Healthy individuals of any age or physical activity level. |
| **Intervention:** | Original studies investigating the effects of oral BA supplementation on skeletal MCarn content. |
| **Comparator:** | No human comparators were required in the studies included in this review, although the data from placebo groups were used to quantify biological variability across the times periods investigated, when available. |
| **Outcomes:** | The primary outcome was the effect of BA supplementation on skeletal MCarn concentration. Potential moderators to this response included dose, sex, age, baseline MCarn and the method used to measure MCarn. |
| **Study Design:** | Controlled or uncontrolled intervention studies. |

### Search Strategy

The search strategy was based on a three-step screening (title/abstract screening, full-text screen and full text appraisal), independently undertaken by two reviewers. This search was originally conducted to inform a systematic risk assessment on the use of BA supplementation [16]. This risk assessment included all BA supplementation studies (including both human and animal models). One hundred and one human studies were included in that investigation, and were subsequently screened to identify those that included a MCarn measurement. The search strategy, including databases and keywords used has been described in detail elsewhere [16] and the protocol for that review was prospectively registered ((PROSPERO registration no. CRD42017071843).

### Data Analysis

The present study comprised both individual and aggregate data meta-analyses from a Bayesian perspective. Individual data were pooled using Bayesian mixed effects multilevel models. Analyses were performed on the outcome variable MCarn (absolute value) to quantify the effects of beta-alanine supplementation and random noise due to biological variation and measurement error. Additionally, proportion of response was estimated across controlled studies by calculating interindividual difference in response to supplementation and comparing this to a non-zero increase in MCarn [17]. Bayesian estimates of the standard deviation in observed change from active and placebo groups were used to obtain the intervention response standard deviation 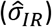 describing interindividual difference in response. Aggregate data meta-analyses were performed using published pre- and post-intervention mean and standard deviation values. Values were transformed into standardized mean differences (SMD) and sampling variance calculated using methods described previously [18]. Three-level mixed effects models were used to quantify the effects of supplementation dose. Insufficient data were available to allow investigation of the interaction between daily dose and intervention duration and so the total cumulative dose ingested was selected as the primary outcome, which previous research has identified as being more influential than either daily dose or intervention duration [16,19,20]. Subset analyses using study covariates were used to assess the effects of sex, age or measurement method on the main effect of BA supplementation. Finally, a model-based approach was employed to investigate the dose-response relationship between cumulative BA supplementation and the SMD. A standard four parameter sigmoid predicted maximum effect (Emax) model was estimated with:

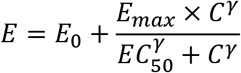

Where *E* is the effect size (SMD), *E*_0_ is the baseline effect, *E_max_* is the maximum effect, *EC*_50_ is the cumulative dose that provides 50% of the maximum effect, *C* is the input (cumulative dose) and *γ* is the Hill coefficient controlling the slope of the sigmoid response. Inferences from all models were performed on posterior samples generated by Markov Chain Monte Carlo with Bayesian 95% credible intervals (CrIs) constructed to enable probabilistic interpretations of parameter values. Models were run in OpenBUGS (version 3.2.3, MRC Biostatistics Unit) and in R (version 3.3.1 R Development Core Team) using the R2OpenBugs package.

## RESULTS

### Study characteristics

Twenty-five studies were identified in the systematic search and included in the meta-analysis [10–12,19–40], along with three, currently unpublished, data sets from the authors’ lab (see Figure 1 for search flow diagram). In total, 575 participants (comprising 486 men and 89 women) were included, of which 382 consumed BA, with the remaining 193 allocated to a placebo intervention. The majority of studies were conducted on healthy young adults (mean age (yrs) = 23.89, SD = 5.46), with only one study conducted on elderly (mean age (yrs) = 64.34, SD = 4.99, [26]). An overview of all included studies is presented in Supplementary File 1. Analyses were completed on subsets of the data depending on the specific analysis and suitability of each study set, as described below.

**Figure 1:**
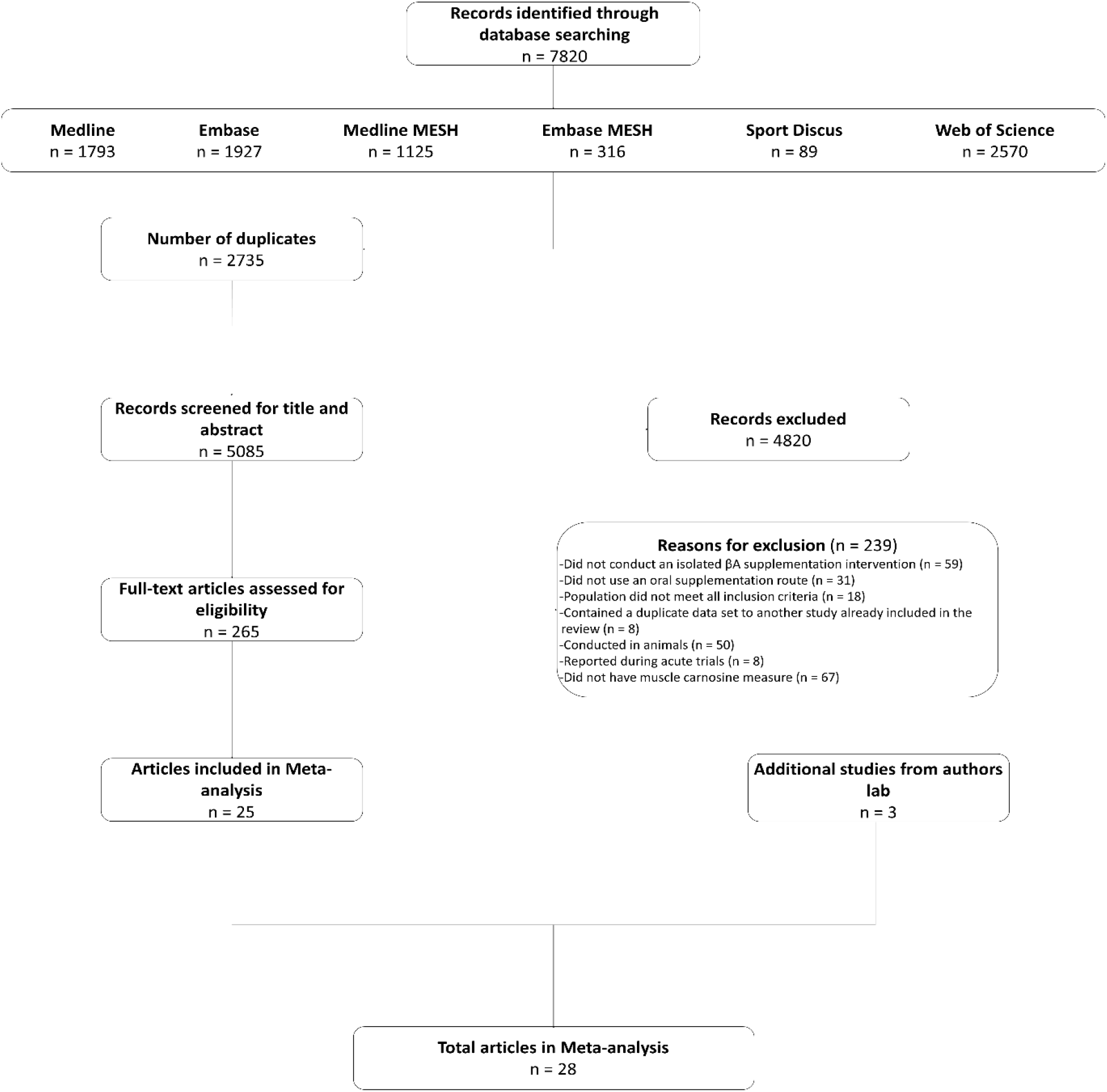
Search Flow Diagram

#### Individual Data

Complete individual data sets were obtained from five studies conducted within the authors laboratory. Two of these studies were identified in the systematic search [12,22], while the remaining three are currently unpublished. Ninety-nine participants were available (BA_n_ = 67, PLA_n_ = 32) comprising a total of 232 observations. All studies provided a BA dose of 6.4 g·day^-1^ with observations ranging from 4 to 24 weeks post baseline. BA supplementation increased MCarn on average by 16.0 mmol·kgDM^-1^ ([95%CrI: 12.4 – 19.6] compared to placebo. Regression analyses with duration centred at 4 weeks were completed to determine if the effects of supplementation increased beyond this point (BA_n_ = 50, 134 total observations). The mean change in MCarn at 4 weeks was 14.0 mmol·kgDM^-1^ [95%CrI: 10.1 – 18.1], with a positive regression slope indicating a further 0.5 [95%CrI: 0.2 – 0.7] mmol·kgDM^-1^ increase per week. Analyses of the same data also demonstrated that baseline levels of MCarn were not associated with changes due to supplementation (−0.1 [95%CrI: −0.3 – 0.1]). The amount of random noise in MCarn values due to biological variation and measurement error (*i.e*., typical variation) was estimated using observations from placebo groups. The standard deviation of residuals from the multilevel model representing typical variation was 4.1 mmol·kgDM^-1^([95%CrI: 3.4-5.1], PLA_n_ = 18, 61 total observations). Calculation of proportion of response first required an estimate of the intervention response standard deviation 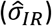, which determines the variability of individual change centered on the group pre to post change. The intervention response standard deviation 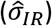 was estimated as 6.6 mmol·kgDM^-1^ [95%CrI: 3.4 – 9.4] and the proportion of response was 99.3% [95%CrI: 96.2 - 100].

#### Aggregate Data

Aggregate analyses were based on effect sizes calculated from all available studies using the SMD pre to post change in MCarn levels. One hundred and eight effect sizes were available from BA groups only, 6 of which were removed as they were outliers (ES > 5). The multilevel meta-analysis with no study covariates estimated a large pooled effect size of 1.5 [95%CrI 1.2 – 1.8], with substantial between (*τ*^2^_0.5_ = 0.6) and within (*ϵ*^2^_0.5_ = 0.7) study variance (Figure 2). The same model applied to effect sizes calculated with supplementation and control group data (22 studies and 56 effect sizes) also produced a large pooled effect size of 1.7 [95%CrI: 1.3 – 2.1], with substantial between (*τ*^2^_0.5_ = 0.8) and within (*ϵ*^2^_0.5_ = 0.5) study variance (Figure 3). Using a simple linear model, the effects of cumulative BA dose was assessed by centering on the mean value (208g). Results demonstrated a large effect at the mean cumulative dose (1.5 [95%CrI: 1.2 – 1.8]) and an estimated 0.23 [95%CrI: 0.06 – 0.49] increase in effect size per additional 100g. Similar results were obtained for effect sizes calculated with supplementation and control group data (effect at mean: 1.7 [95%CrI: 1.3 – 2.1]; effect per additional 100g: 0.16 [95%CrI: 0.01 – 0.31]). Insufficient data were available to ascertain if age altered the effects of BA supplementation, but subset analyses were conducted to investigate the impact of sex and the method used to measure MCarn, using effect sizes generated from supplementation groups only. Sixteen studies were selected that used the most common dosing protocol (cumulative dose between 130 and 180g) comprising a total of 56 effect sizes. For the sex comparison there were 8 effect sizes from a female only group, 38 effect sizes from a male only group and 10 effect sizes from a mixed group. No substantive evidence of a gender effect was obtained (male vs female: −0.32 [95%CrI:-1.1 – 0.43]; male vs mixed: −0.00 [95%CrI: −0.95 – 0.88]). Across the 16 studies, 40 effect sizes were obtained from MCarn values measured with non-invasive scanning devices (*i.e*., HR-MRS) and 16 effect sizes obtained with muscle biopsy based analyses (mainly assessed by HPLC, with one study using UPLC and one using mass spectrometry), with some evidence of increased effects with HPLC (0.16 [95%CrI: 0.01 – 0.43]).

**Figure 2:**
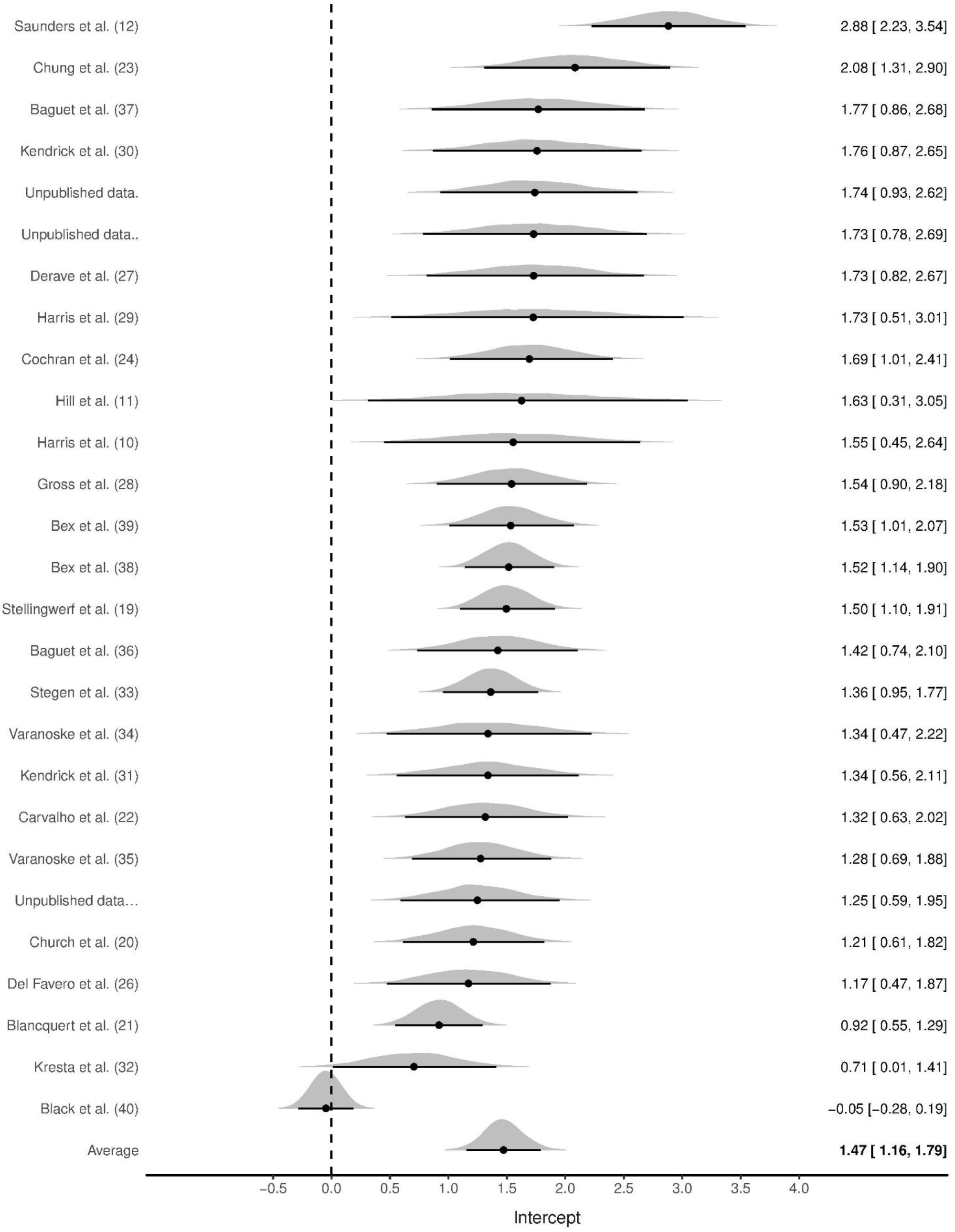
Bayesian Forest Plot of multilevel meta-analysis with non-controlled effect sizes

**Figure 3:**
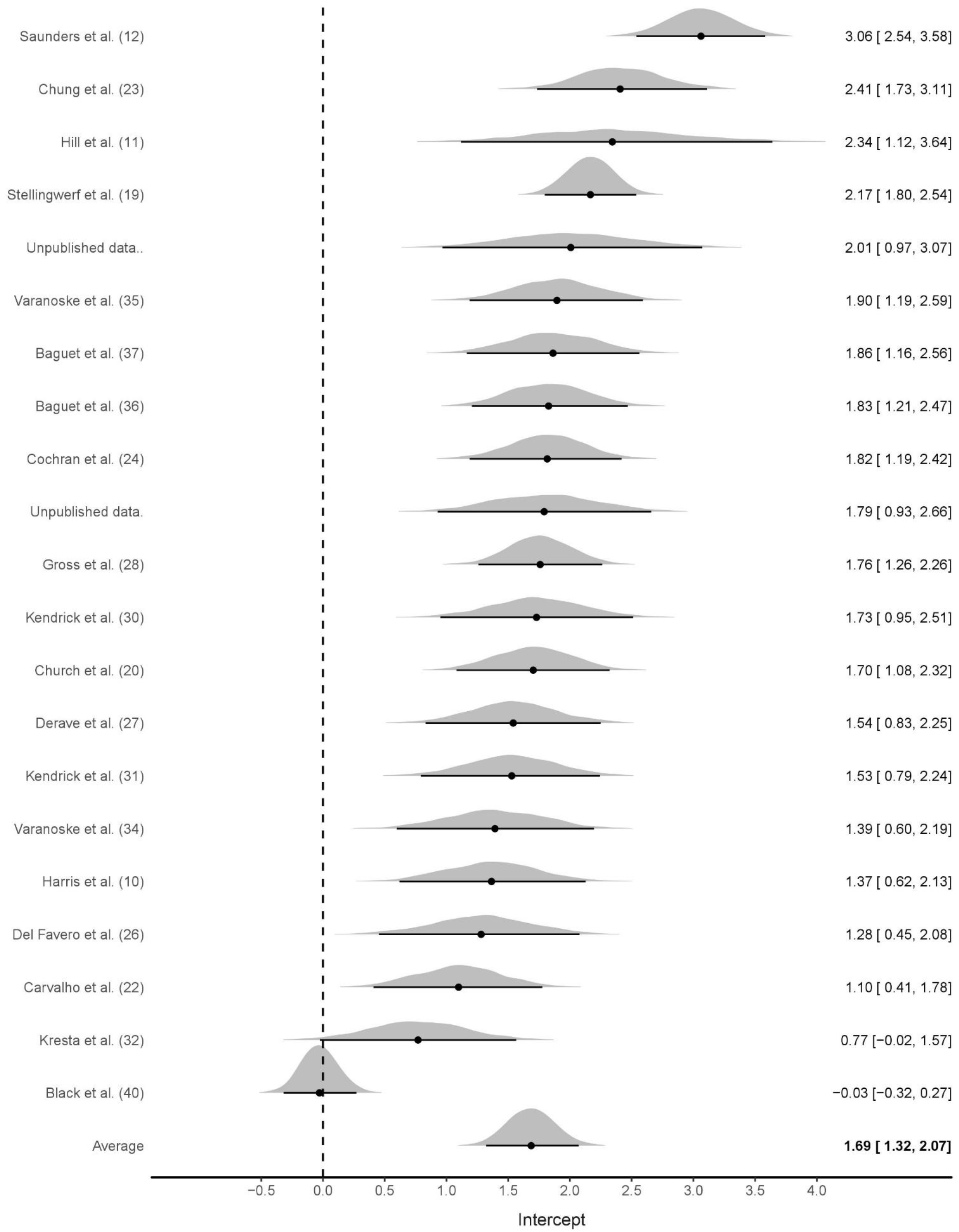
Bayesian Forest Plot of multilevel meta-analysis with controlled effect sizes

#### Emax Model

The predicted maximum effect of BA supplementation (Emax) was 3.0 (50%CrI 2.2 – 3.7) and the estimated total cumulative dose (g) required to achieve 50% of this maximum effect (ED50) was 377g (50%CrI 210 – 494). A density plot with the Emax curve generated from median parameter values is provided in Figure 4. An extrapolation of posterior samples from the Emax model was performed to estimate probabilities that percentage of maximum effect could be achieved with cumulative doses ranging from 1000 to 1500g (see Table 2). These results estimated, for example, that the probability of obtaining at least 70% of maximum effect with a cumulative dose of 1000g was 0.68.

**Figure 4:**
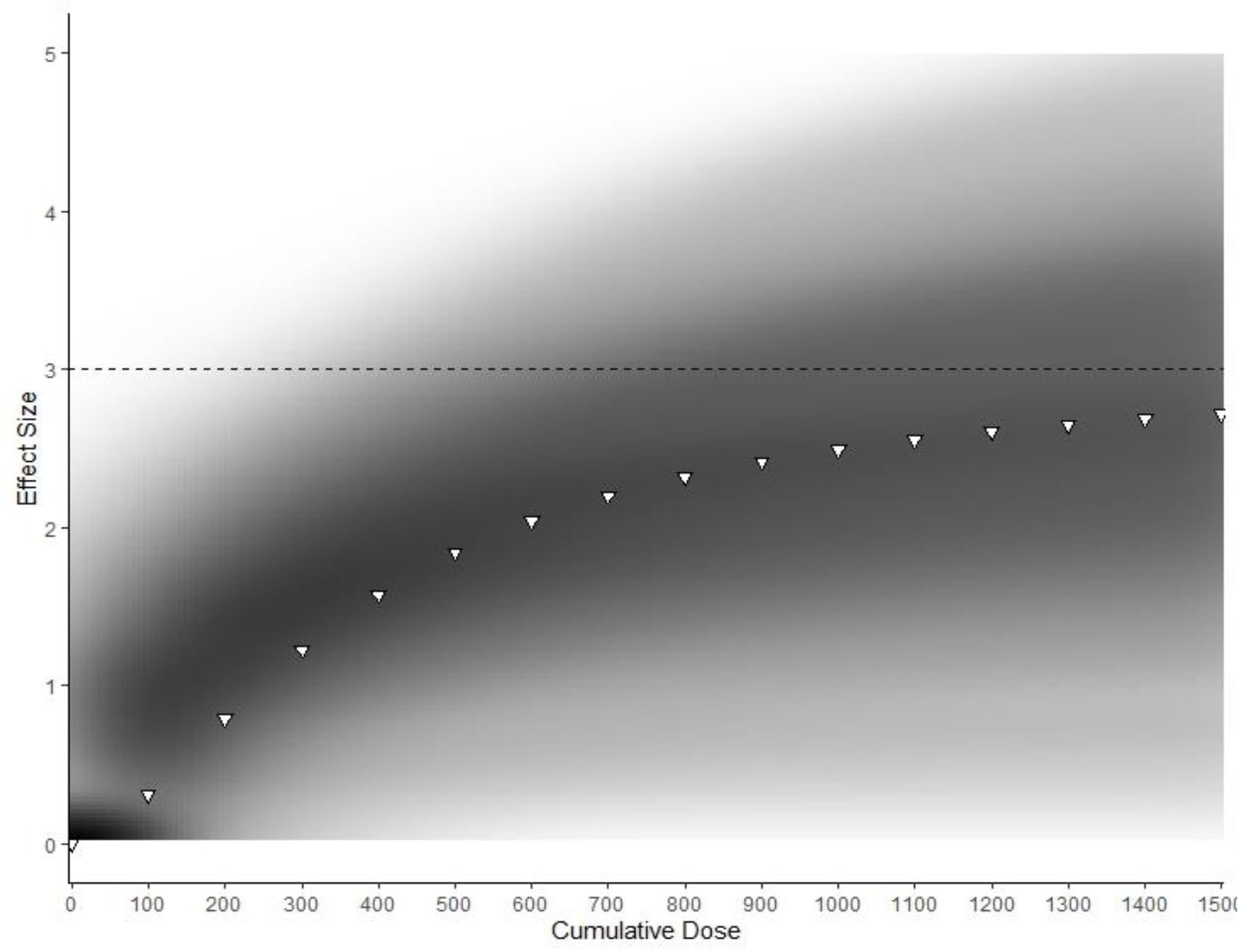
Density plot of Bayesian Emax model predicting effect of cumulative BA supplementation on muscle carnosine content. ***<u>Note:</u>** Darker areas represent more common Emax trajectories. White triangles represent Emax generated with median parameter values*.

**Table 2:** Probability table representing the chance that various cumulative doses (columns) create a response greater than the specified percentage of EMax (rows) based on Bayesian model generated.

| Percentage of E<sub>max</sub> | 1000g | 1100g | 1200g | 1300g | 1400g | 1500g |
| --- | --- | --- | --- | --- | --- | --- |
| 70% | 0.68 | 0.73 | 0.77 | 0.80 | 0.83 | 0.85 |
| 80% | 0.45 | 0.48 | 0.51 | 0.54 | 0.56 | 0.59 |
| 90% | 0.31 | 0.33 | 0.35 | 0.37 | 0.38 | 0.40 |

Values in table represent probabilities (p) 0 ≤ p ≤ 1.

## Discussion

The purpose of this study was to conduct a comprehensive analysis with various modelling techniques to synthesize existing knowledge about the MCarn response to BA supplementation. Collectively, our findings and models employed indicated that human skeletal muscle has large capacity for MCarn accumulation, and that commonly used protocols (*e.g*., 4 weeks at 6.4 g·day^-1^) may not come close to saturating muscle carnosine content. Baseline values do not appear to influence subsequent response to supplementation and the non-linear response to supplementation was not influenced by sex. Analysis of individual data indicated that muscle carnosine content is relatively stable in the absence of the intervention, and that effectually all (99.3% [95%CrI: 96.2 – 100]) participants respond to BA supplementation.

Our observation that humans have large capacity for non-linear MCarn accumulation align with a recent theoretical model proposed by Spelnikov & Harris [41]. Their model describes changes in MCarn over time with BA supplementation, and was based on three studies that used a slow release BA formulation. The authors described absolute increases in MCarn as a product of the rates of synthesis and decay, with carnosine synthesis considered to be constant in relation to time and first order to daily BA dose. Similarly, carnosine decay was also considered to be first order, but in relation to total MCarn content. As such, carnosine decay increases when absolute content is higher and so the rate of MCarn accumulation due to BA induced elevations in synthesis will slow. Tissue saturation represents the point at which the rates of synthesis match decay, and so content will remain constant despite continued supplementation. The lack of data at higher BA doses limits precision regarding the point at which human skeletal muscle saturation may occur; however, our analyses suggest that humans have very large capacity to accumulate MCarn and that naturally occurring baseline levels are far below that which the muscle is capable of maintaining.

Our analyses indicate that the nature of individual MCarn response to BA supplementation differs from other commonly used supplements, such as creatine. Human skeletal muscle appears to reach creatine saturation at approximately 140 – 160 mmol·kgDM^-1^ [42] and this can be achieved within 5 days of high-dose supplementation. Response to creatine supplementation is largest in those with lowest baseline levels, whereas individuals whose creatine content is habitually closer to this saturation point gain smaller benefit from supplementation [42]. In contrast, we observed no evidence of an influence of baseline carnosine content on the subsequent response to supplementation. This makes sense when considered in relation to our predictive model, as it appears that we have very large capacity to accumulate MCarn – far greater than is achieved with commonly used protocols (*e.g*., 179.2 grams provided as 6.4g·day^-1^ for 4 weeks). This may be because baseline MCarn contents (approximately 25 mmol·kgDM^-1^) are substantially lower than those observed for creatine, with many individuals having baseline contents close to the proposed creatine saturation limit of 140 – 160 mmol·kgDM’^1^. These data, in turn, raise another interesting question. Does human skeletal muscle have a largely uniform saturation point, after which no further increases can be attained (as seems to be the case with creatine)? Or does capacity to accumulate MCarn vary widely between individuals, with each having their own upper limit? Currently, insufficient data using very high BA protocols on MCarn precludes the answering of this question, but one thing that is clear is that human skeletal muscle has large capacity to uptake BA and to increase MCarn above naturally occurring, non-supplemented levels.

Our data indicated that humans have large capacity for MCarn accumulation, and this, in turn, raises other important questions, *e.g*., is maximal MCarn accumulation necessary, or even desirable? Theoretically, the greater the increase in MCarn content, the greater its ability to buffer, and to contribute to other processes, and so intuitively, attaining the largest increases possible seems desirable. Two studies did report that larger MCarn increases were associated with greater performance effects [11,12], but meta-analytic data does not support this, and the total dose ingested was previously reported not to influence the effect on exercise performance [18]. It would be counterintuitive to believe that performance benefits would continue to linearly increase with ever-increasing MCarn, given that numerous factors, apart from acidosis, contribute to fatigue [43], and so it makes sense that at some point, performance benefits would plateau. Numerous studies have reported that approximately 179g of BA can be ergogenic, and so it seems MCarn saturation is not essential for BA supplementation to be ergogenic, although it does remain to be seen whether greater increases can elicit greater effects on exercise performance. A very important avenue for future research is the identification of the lowest MCarn increase necessary to elicit an ergogenic effect, along with the point after which no further benefits can be obtained. This information could have large potential to enhance the applicability and efficacy of BA supplementation strategies. For example, it seems that the largest gains in MCarn are attained in the earlier phases of supplementation (see Figure 4). It would be of interest to identify if strategies such as meal co-ingestion [33], intake in proximity to training [39] or intake in slow-release capsules [35] can influence the early response to supplementation [14] and whether this, in turn, meaningfully impacts exercise performance.

The current analysis also brought to light some interesting points about the nature of carnosine response to supplementation, which has implications for future study design. It seems that in the absence of intervention carnosine is relatively stable in the muscle, likely due to low intramuscular carnosinase and roughly equivalent synthesis and degradation rates [3]. Our analysis of individual data indicated typical variation of approximately 4 mmol·kgDM^-1^ across a 4-week period. Previously, two muscle samples taken from the same biopsy cut showed a variation of approximately 1mmol·kgDM^-1^ [44], and so measurement error likely accounts for at least a quarter of this variation (and potentially more). It was interesting to observe that both within and between study variance were large and similar. A large proportion of this sampling error is likely due to small sample sizes. Typically the use of a control group would be recommended to normalize the effects of the intervention against those of usual biological variability [17]. But in this situation we observed little variation in placebo group MCarn, while the effects of intervention studies when analysed both with and without controlling for the effects of the placebo group were similar (ES (95%CrI): 1.7 (1.3 – 2.1) versus 1.5 (1.2 – 1.8). This implies that the control group adds little value to the analysis, likely because of MCarn stability and the large effect of supplementation. In future investigations of the MCarn response to BA in young healthy males (and particularly those for which resources are limited) it may be prudent to direct resources toward the intervention group, in order to reduce within study variance. It is important to note that this recommendation applies only to studies on the MCarn response to BA supplementation. The influence of BA supplementation on exercise performance is far less well-characterized and subject to substantially more sources of internal and external variability and so control groups are essential in studies for which exercise is the primary outcome of interest.

In addition to characterizing the nature of MCarn response to BA supplementation, we also considered the influence of various potential moderators on this response. In relation to the method of assessment, it seems that lower effect estimates are generally observed when MCarn is measured using the HR-MRS technique when compared to those obtained using HPLC analysis of muscle biopsies. When considering the influence of non-modifiable factors on the MCarn response to supplementation (namely age and sex), we were unable to conduct analyses on the influence of age, as most studies were conducted in young men, and insufficient data in older and younger groups was available. Women have previously been reported to have lower MCarn than men [45,46], but our data indicate that both men and women have a similar response to BA supplementation. This implies that the lower values previously reported in women are unlikely to relate to an inherent gender dysmorphism in the biological factors that underpin carnosine metabolism.

In conclusion, our findings indicate that human skeletal muscle has large capacity to accumulate carnosine. MCarn remains stable in the absence of intervention and neither low baseline MCarn levels, nor sex, influence the subsequent response to BA supplementation. In turn, these findings lead to other questions, the response to which may have large implications for future practice. From the point of view of athletic performance, key questions include: what is the absolute MCarn increase required to elicit an ergogenic effect, along with the point after which no further benefits are attained? It is clear that 4 weeks of BA supplementation can be ergogenic, but can this be achieved earlier? Can strategies to enhance the early response to BA supplementation meaningfully impact the subsequent ergogenic benefits? The response to these questions may progress practical application of this supplementation strategy, with potential benefit to many athletic and clinical populations.

